# Integrated glycoproteomics identifies a role of *N*-glycosylation and galectin-1 on myogenesis and muscle development

**DOI:** 10.1101/2020.06.29.178772

**Authors:** Ronnie Blazev, Christopher Ashwood, Jodie L. Abrahams, Long H. Chung, Deanne Francis, Pengyi Yang, Kevin I. Watt, Hongwei Qian, Gregory A. Quaife-Ryan, James E. Hudson, Paul Gregorevic, Morten Thaysen-Andersen, Benjamin L. Parker

## Abstract

Many cell surface and secreted proteins are modified by the covalent addition of glycans that play an important role in the development of multicellular organisms. These glycan modifications enable communication between cells and the extracellular matrix via interactions with specific glycan-binding lectins and the regulation of receptor-mediated signaling. Aberrant protein glycosylation has been associated with the development of several muscular diseases suggesting essential glycan- and lectin-mediated functions in myogenesis and muscle development but our molecular understanding of the precise glycans, catalytic enzymes and lectins involved remain only partially understood. Here, we quantified dynamic remodeling of the membrane-associated proteome during a time-course of myogenesis in cell culture. We observed wide-spread changes in the abundance of several important lectins and enzymes facilitating glycan biosynthesis. Glycomics-based quantification of released *N*-linked glycans confirmed remodeling of the glycome consistent with the regulation of glycosyltransferases and glycosidases responsible for their formation including a previously unknown di-galactose-to-sialic acid switch supporting a functional role of these glycoepitopes in myogenesis. Furthermore, dynamic quantitative glycoproteomic analysis with multiplexed stable isotope labelling and analysis of enriched glycopeptides with multiple fragmentation approaches identified glycoproteins modified by these regulated glycans including several integrins and growth factor receptors. Myogenesis was also associated with the regulation of several lectins most notably the up-regulation of galectin-1 (LGALS1). CRISPR/Cas9-mediated deletion of *Lgals1* inhibited differentiation and myotube formation suggesting an early functional role of galectin-1 in the myogenic program. Importantly, similar changes in *N*-glycosylation and the up-regulation of galectin-1 during postnatal skeletal muscle development were observed in mice. Treatment of new-born mice with recombinant adeno-associated viruses to overexpress galectin-1 in the musculature resulted in enhanced muscle mass. Our data form a valuable resource to further understand the glycobiology of myogenesis and will aid the development of intervention strategies to promote healthy muscle development or regeneration.

## INTRODUCTION

The bulk of skeletal muscle is composed of post-mitotic multi-nucleated myofibers that form via the fusion of mono-nucleated progenitor myoblasts. Myofiber formation is achieved via myogenesis, a highly ordered process including differentiation, elongation, migration, cell adhesion, membrane alignment and ultimately cell fusion of myoblasts (1). The initial differentiation of myoblasts is regulated by external growth factors, cytokines, steroid hormones and signal transduction pathways that activate a series of muscle-specific and pleiotropic transcription factors (2). Elongation of myoblasts is achieved by extension of filopodia and lamellipodia to contact surrounding muscle cells. Myoblasts subsequently migrate to each other, which requires extracellular matrix (ECM) remodeling to facilitate cell motility prior to cell recognition and adherence. Here, interactions between multiple adherence molecules trigger integrin signaling and a re-arrangement of the actin-cytoskeleton. This is coupled to the regulation of several GTPases and guanine nucleotide exchange factors that contribute to membrane remodeling and cell fusion via the ARP2/3, WASP and WAVE protein complexes (3). In mammals, distinct phases of myogenesis contribute to the formation of mature skeletal muscle (4, 5). Muscle patterning is established by the fusion of embryonic myoblasts. The second phase involves fusion of fetal myoblasts followed by the formation of the basal lamina and the expansion of adult precursor satellite cells (muscle stem cells). Accompanying this second phase is innervation of the myofibers leading to the formation of the neuromuscular junctions (NMJ). Finally, postnatal myogenesis is achieved via myoblasts derived from satellite cells that are responsible for growth and regeneration of mature skeletal muscle.

Myogenesis and muscle development involve the interaction of cell surfaces and ECM with hundreds of glycosylated proteins. It is therefore not surprising that defects in glycosylation have been associated with several developmental disorders. More than 50 congenital disorders of glycosylation (CDGs) have been identified in humans, and these typically present as abnormalities in development of the nervous system and/or skeletal muscle during infancy (6). The majority of CDGs are inherited defects in one or more of the multiple enzymes responsible for glycosylation of asparagine residues (*N*-linked glycosylation) that occur on membrane-associated, cell surface and secreted proteins. For example, several loss-of-function mutations have been identified in the *PMM2* gene involved in the synthesis of GDP-mannose, a nucleotide-sugar donor responsible for the transfer of mannose residues to *N*-glycans and thus critical for their maturation. Individuals with PMM2-CDG have hypo-*N*-glycosylation, and display severe hypotonic muscles and underdeveloped cerebellum (7). Advances in next generation DNA sequencing have pin-pointed additional CDGs in patients presenting with abnormal *N*-glycosylation and defects in muscle and/or nervous system development. This includes the identification of mutations in the *STT3A* and *STT3B* genes forming the catalytic subunits of the N-oligosaccharyl transferase (OST) complex (8), and *MAN1B1*, which are involved in the regulation of *N*-linked glycosylation (9). Despite the well documented mutations in genes regulating glycosylation and their associations with poor muscle function, we know very little about the regulation of *N*-glycosylation during myogenesis and muscle development. Furthermore, the roles of glycan-binding proteins (GBPs) such as lectins on myogenesis and muscle development remain only partially understood. For example, loss of *Lgals1* expression results in defects in muscle development in mice and fish (10, 11). The molecular mechanisms remain poorly defined but *in vitro* experiments suggest Galectin-1 binds to a variety of ECM proteins including laminins and integrins at the sarcolemma in order to regulate intracellular signaling (12). Muscular dystrophies are characterized by progressive muscle degeneration, sarcolemmal damage and loss of muscle function. Analysis of immortalized healthy and dystrophic human muscle cells with lectin histochemistry has recently revealed several changes in lectin binding suggesting discrete changes in glycosylation or glycoprotein abundance (13). Excitingly, administration of recombinant galectin-1 in a genetic mouse model of muscular dystrophy (new born *mdx* mice) display improved muscle function and sarcolemmal integrity suggesting critical roles of galectin-1 in myogenesis (14). These experiments are encouraging as it highlights new potential treatment options for a variety of muscle diseases. However, further experiments are required to characterize the role of *N*-glycosylation and galectin-1 on myogenesis and muscle development.

## EXPERIMENTAL PROCEDURES

### L6 cell culture

Rat L6 myoblasts were maintained in α-minimum essential medium (α-MEM) containing 5.5 mM glucose (Gibco) and 10% fetal bovine serum (FBS; Hyclone Laboratories) in a 10% CO2 incubator. L6 myoblasts were differentiated into myotubes over 7 days with 2% FBS when myoblasts reached ~90-95% confluency. For CRISPR/Cas9 experiments targeting complete disruption of *Lgals1*, myoblasts were transfected at 60% confluency with DNA constructs expressing CMV-Cas9(D10A) and paired U6-gRNAs (5’-GTTGTTGCTGTCTTTCCCCAGG and 5’- ACCCCCGCTTCAACGCCCATGG) (Sigma Aldrich) using TransIT-X2 reagent (Mirus Bio). After 48 hours, cells were trypsinized, counted and serial-diluted to ~0.8 cells per 30 μl in fresh conditioned α-MEM media containing 10% FBS. Cells were seeded into 384-well plates with media replaced every 2-3 days and single clones selected. After 10 days, cells were expanded and screened for LGALS1 ablation using western-blot analysis.

### L6 cell lysis for mass spectrometry analysis

Cells were washed twice with ice-cold PBS and lysis in ice-cold 100 mM sodium carbonate containing protease inhibitor cocktail (Roche) by tip-probe sonication. Lysates were rotated at 4℃ for 1 h then centrifuged at 150,000 x g for 60 min at 4℃ to pellet microsomal protein fraction. The pellet was resuspended in 6 M urea, 2M thiourea, 1% SDS containing 25 mM triethylammonium bicarbonate (TEAB), pH 8.0 and protein precipitated with chloroform:methanol:water (1:3:4). Precipitated protein was washed with methanol and resuspended in 6 M urea, 2 M thiourea containing 25 mM TEAB, pH 8.0. Protein concentration was determined via Qubit (Invitrogen), normalized and stored at −80℃.

### (Glyco)proteomic sample preparation

Peptides were prepared essentially as described previously (15). Briefly, 35 μg of membrane-enriched protein from each of the eight time points (0-7 days) and each biological replicate (n = 3) were reduced with 10 mM dithiothreitol (DTT) for 1 h at room temperature followed by alkylation with 25 mM iodoacetamide for 30 min at room temperature in the dark. The reaction was quenched to 25 mM DTT and digested with 0.7 μg of sequencing-grade LysC (Wako Chemicals) for 3 h at room temperature. The digest was diluted 5-fold with 25 mM TEAB and digested with 0.7 ug of sequencing-grade trypsin (Sigma) overnight at 30℃. Samples were acidified to a final concentration of 2% formic acid (FA), centrifuged at 13,000 x g for 10 min at room temperature and desalted using 30 mg of hydrophilic-lipophilic balance – solid phase extraction (HLB-SPE) material in a 96-well plate (Waters) using a vacuum manifold. The plate was washed with 5% acetonitrile (MeCN) containing 0.1% trifluoroacetic acid (TFA). The peptides were eluted with 50% MeCN containing 0.1% TFA and dried by vacuum centrifugation. Peptides were resuspended in 20 μl of 100 mM HEPEs, pH 8.0 and isotopically labelled with 73 μg of 10-plex tandem mass tags (TMT) in a final concentration of 33% MeCN for 2 h at room temperature. The sample channels were labelled as follows: 126 = day-0, 127N = day-1, 127C = day-2, 128N = day-3, 128C = day-4, 129N = day-5, 129C = day-6, 130N = day-7. The reactions were quenched with 0.8% hydroxylamine for 15 min at room temperature, and then acidified and diluted to a final concentration of 0.1% TFA and 5% MeCN. The labelled peptides were pooled and purified by HLB-SPE as described above. Peptides were resuspended in 90% MeCN containing 0.1% TFA and 10% of the peptide separated directly into 12 fractions for total proteome analysis using amide-80 HILIC as described previously (16). Glycopeptides were enriched from 90% of the remaining peptide using zic-HILIC microcolumns as previously described (17, 18).

### (Glyco)proteomic mass spectrometry

The analysis of each fraction for proteome quantification was performed on a Dionex 3500RS coupled to an Orbitrap Fusion (Thermo Scientific) operating in positive polarity mode. Peptides were separated using an in-house packed 75 μm x 50 cm pulled column (1.9 μm particle size, C18AQ; Dr Maisch, Germany) with a gradient of 2 – 40% MeCN containing 0.1% FA over 150 min at 250 nl/min at 55°C. MS1 scans were acquired from 350 – 1,400 *m/z* (120,000 resolution, 5e5 AGC, 50 ms injection time) followed by MS/MS data-dependent acquisition of the 10 most intense ions with CID and detection in the ion-trap (rapid scan rate, 1e4 AGC, 70 ms injection time, 30% NCE, 1.6 *m/z* isolation width). Synchronous precursor selection was enabled with multi-notch isolation of the 10 most abundant fragment ions excluding precursor window of 40 *m/z* and loss of TMT reporter ions for MS3 analysis by HCD and detection in the Orbitrap (60,000 resolution, 1e5 AGC, 300 ms injection time, 100 – 500 *m/z*, 55% NCE, 2 *m/z*) (19).

The analysis of glycopeptides was performed as single-shot analysis without fractionation on the identical system as described above. MS1 scans were acquired from 550 – 1,750 *m/z* (120,000 resolution, 5e5 AGC, 100 ms injection time) followed by MS/MS data-dependent acquisition of the 7 most intense ions and highest charge state with HCD and detection in the Orbitrap (60,000 resolution, 2e5 AGC, 200 ms injection time, 40 NCE, 2 *m/z* quadrupole isolation width). The acquisition strategy included a product ion triggered re-isolation of the precursor ion if HexNAc oxonium ions (138.0545 and 204.0867 *m/z*) were detected amongst the top 20 fragment ions of the HCD-MS/MS spectrum. The re-isolated precursor ions were subjected to both EThcD- and CID-MS/MS analysis (20–22). EThcD-MS/MS analysis was detected in the Orbitrap (60,000 resolution, 3e5 AGC, 250 ms injection time, calibrated charge dependent ETD reaction times [2+ 121 ms; 3+ 54 ms; 4+ 30 ms; 5+ 20ms; 6+ 13 ms; 7+; 10 ms], 2 m/z quadrupole isolation width) and, CID MS/MS analysis was detected in the ion trap (rapid scan rate, 1e4 AGC, 70 ms injection time, 35% NCE, 2 *m/z* quadrupole isolation width). Data are available via ProteomeXchange with identifier PXD019372 (23). Username: and Password: KGAgOAkh

### (Glyco)proteomic data analysis

The identification and quantification of peptides for proteomic analysis was performed with Proteome Discoverer (v2.1.0.801) using Sequest (24) against the rat UniProtKB database (November 2015). The precursor mass tolerance was set to 20 ppm with a maximum of two full trypsin miss cleavages while the CID-MS/MS fragment mass tolerance was set to 0.6 Da. The peptides were searched with oxidation of methionine set as variable modifications while carbamidomethylation of cysteine and TMT of peptide N-terminus and lysine was set as a fixed modification. All data were searched as a single batch with PSM and peptide FDR filtered to 1% using Percolator (25) and protein level FDR set to 1% using Protein FDR Validator node. Quantification was performed using the Reporter Ions Quantifier node with integration set to 20 ppm and co-isolation threshold set to 50% and reporter ions were required in all channels. Peptides were grouped in each replicate based on unique sequence and unique modifications, and the median reporter ion areas calculated. Data were expressed as Log2 fold-change to day-0 for each replicate and normalized to a median of 0. Total proteome from biological replicates across time points were batch effect corrected using an empirical Bayes model implemented in the sva R package (26). As recommended, the parametric shrinkage adjustment was applied to the data. The quality of the data after batch effect correction was assessed using the principal component analysis (PCA) and the corrected data were used for subsequent analysis. Significantly regulated glycopeptides were determined using ANOVA with permutation-based FDR set at 5% with Tukey’s posthoc test.

The identification and quantification of glycopeptides was performed with Proteome Discoverer (v2.1.0.801) using the Byonic node (27) against the rat UniProtKB database (November 2015). The precursor, HCD- and EThcD-MS/MS mass tolerance were set to 20 ppm with a maximum of two full trypsin miss cleavages. The peptides were searched with oxidation of methionine and N-glycan modification of asparagine (309 possible glycan compositions without sodium adducts available within Byonic) set as variable modifications while carbamidomethylation of cysteine and TMT of peptide N-terminus and lysine were considered as fixed modifications. A precursor isotope off set was enabled (narrow) to account for incorrect precursor monoisotopic identification (+/− 1.0 Da). All data were searched as a single batch with PSM FDR set to 1% using the PSM Validator node and a minimum Byonic score of 100 was applied (28). Only HCD-MS/MS spectra containing HexNAc oxonium ions (138.05-138.06 and 204.08-204.09 *m/z*) were annotated in the final list of glycopeptides as previously described (29). All identified glycopeptides were quantified based on HCD-MS/MS data using the Reporter Ions Quantifier node with integration set to 20 ppm and co-isolation threshold set to 75%. Reporter ions were required in all channels. Peptides were grouped in each replicate based on unique sequence and unique modifications, and the median reporter ion areas calculated. Data were expressed as Log2 fold-change to day-0 for each replicate and normalized to a median of 0. Significantly regulated glycopeptides were determined using ANOVA with permutation-based FDR set at 5% with Tukey’s posthoc test.

### Glycomic sample preparation

*N*-glycome profiling was performed essentially as described previously (30). Briefly, 10 μg protein extracts were dot-blotted onto PVDF membranes and allowed to dry overnight. The membranes were stained briefly with direct blue 71 in 40% ethanol containing 10% acetic acid and washed with water. Immobilized proteins were excised and the membrane blocked with 1% polyvinyl pyrrolidine 4000 for 5 min followed by washing with water. *N*-glycans were released with 2.5 U PNGase F (Roche, Australia) for 16 h at 37°C. Released glycans were collected, incubated with 100 mM ammonium acetate, pH 5.0 for 1 h at 23°C and dried by vacuum centrifugation. *N*-glycans were reduced with 1 M NaBH_4_ in 50 mM KOH for 3 h at 50°C then desalted and enriched offline using AG 50W-X8 (Bio-Rad, Australia) strong cation exchange followed by porous graphitized carbon (PGC) solid phase extraction micro-columns (Grace, USA). For the determination of galactose linkages, aliquots of released glycans were incubated with combinations of 20 U broad-specificity sialidase (P0722S, α2-3,6,8,9), 8 U broad-specificity α-galactosidase (P0747S, α1- 3,4,6), 10 U β-galactosidase (P0726S, β1-3) and 8 U β-galactosidase (P0746S, β1-3,4) (all from New England BioLabs, USA). All reactions were performed in a final volume of 10 μl in 50 mM sodium acetate buffer, 5 mM CaCl_2_ pH 5.5 for 16 h at 37°C.

### Glycomic mass spectrometry

PGC-LC-ESI-MS/MS experiments were performed on an 3D ion trap using an Agilent 1100 capillary LC system (Agilent Technologies) interfaced with an Agilent 6330 LC-MSD 3D Trap XCT ultra. A PGC LC column (3 μm, 100 mm × 0.18 mm, Hypercarb, Thermo Scientific) was maintained at room temperature and at 50°C. 10 mM ammonium bicarbonate aqueous solution (solvent A) and 10 mM ammonium bicarbonate aqueous solution with 45% MeCN (solvent B) were used as mobile phases at 2 μl/min with the following gradient: 0 min, 2% B; linear increase up to 35% B for 53 min; linear increase up to 100% B for 20 min; held constant for 5 min; and then equilibrated at 2% B for 5 min before the next injection—giving a total LC run time of 83 min. Glycans were analyzed using the following ESI-MS conditions: source voltage −3.2 kV, MS1 scan range 350 – 2200 m/z, 5 microscans, 0.13 *m/z* resolution (FWHM), 8e4 ion current control (ICC) and 200 ms accumulation time. Ion trap CID-MS/MS conditions were as follows: 0.13 *m/z* resolution (FWHM), 8e4 ICC, 200 ms accumulation time, 4 *m/z* isolation width and data-dependent acquisition of the three most abundant glycan precursors in each scan. The CID-MS/MS used ultra-pure helium as the collision cell gas. Fragmentation amplitude was set to 1 V with Smart-Frag enabled ramping from 30 to 200% of the fragmentation amplitude for CID-MS/MS with an activation time of 40 ms.

### Glycomic data analysis

Data were analyzed as described previously (31). Briefly, a candidate list of glycans from relevant spectra of each sample were extracted using the ESI-compass v1.3 Bruker Daltonic Software (Bruker DALTONIK GmbH). The extracted monoisotopic precursor masses were searched against GlycoMod (http://www.expasy.ch/tools/glycomod) to identify putative monosaccharide compositions. Interpretation and validation of glycan identities, assisted by GlycoWorkBench v2.1 (available from https://code.google.com/archive/p/glycoworkbench) were based on the existence of A-, B-, C-, X-, Y-, and Z-product ions consistently found across the majority of the CID-MS/MS scans over the elution times of the respective precursor ions. Glycan annotation nomenclature was presented as described previously (32).

Relative glycan quantitation based on glycan precursor intensity was performed using Skyline. The relative abundance of each glycan was determined by the relative AUC peak areas of the monoisotopic m/z for relevant precursors divided by the AUC peak area sum of all glycans identified. The precursor mass analyzer was set to a quadrupole ion trap (QIT) with resolution of 0.5 m/z and precursor ions within the range 50 – 2,000 *m/z* were included. Data are freely available on Glycopost (33) with the identifier GPST000079 (raw files fhttps://glycopost.glycosmos.org/preview/7030709905ed7e73ea0928, PIN: 5219) and Panorama Public (34) (skyline assays, https://panoramaweb.org/RatMuscleGlyco.url).

### Cell and tissue lysis for western-blot analysis

Cells were washed twice with ice-cold PBS and lysis in 2% SDS by tip-probe sonication while frozen muscle was sonicated directly in 2% SDS. Lysates were then centrifuged at 16,000 x g for 10 min at room temperature. Protein concentrations were determined via the BCA (Thermo Fisher), normalized and diluted with Laemlli Buffer. Samples were incubated at 65℃ and 10 μg of protein was separated by SDS-PAGE using 10% gels. Proteins were transferred to PVDF membrane and subjected to immunoblot analysis with anti-LGALS1 (#12936; Cell Signaling Technologies) and pan 14-3-3 loading control (#8312; Cell Signaling Technologies). Detection was achieved with HRP-conjugated donkey anti-rabbit secondary antibody (Jackson ImmunoResearch) and imaged by the Bio-Rad ChemiDoc Gel Imager system.

### Immunofluorescence microscopy

Cells were grown on Matrigel coverslips and analyzed 3 days post-differentiation. Coverslips were washed twice with ice-cold PBS and fixed with 4% paraformaldehyde in PBS for 10 min at room temperature. The cells were washed three times with PBS and permeabilized in 0.2% triton X-100 in PBS for 10 min at room temperature. Three additional washes were performed and then blocked with 3% BSA in PBS for 1 h at room temperature. Cells were stained with MY-32 anti-myosin primary antibody (M4276; Sigma) at 1:400 dilution in PBS containing 1% BSA overnight at 4℃. The cover slips were washed tree times with PBS and stained with goat anti-mouse secondary antibody Alexa Fluor 488 (A28175; Invitrogen) at 1:500 dilution in PBS containing 1% BSA for 1 h at room temperature. The cells were washed three times and stained with Hoechst 33342 (62249; ThermoFisher Scientific) at 1 μg/ml in PBS for 5 min at room temperature. The coverslips were washed three times and mounted on glass slides and imaged on a Nikon C2 confocal microscope. Data was processed in ImageJ using Analyze Particle function to count nuclei (35).

### Generation of recombinant adeno-associated viral serotype-6 (rAAV6) vectors

DNA constructs expressing mouse *Lgals1* were sub-cloned into AAV:MCS-SV40pA plasmid (VectorBuilder). Recombinant adeno-associated viral vectors were generated by co-transfection of 10 μg of plasmids containing cDNA constructs with 20 μg of pDGM6 packaging plasmid into HEK293 cells (seeded 16 h prior at a density equivalent to 3.2-3.8 × 10^6^ cells per P-100 tissues culture plate) using the calcium phosphate precipitate method to generate type-6 pseduotyped viral vectors as previously described (36). After 72 h, cells and culture media were collected, subjected to cell lysis by freeze thawing, and clarified using a 0.22 μm filter (EMD Millipore). Virus was purified by heparin affinity chromatography on an Akta FPLC (HiTrap, GE Healthcare), ultra-centrifuged overnight and resuspended in sterile Ringer’s solution. Vector concentrations were determined using qPCR (Applied Biosystems).

### Intramuscular injections of AAV6 vectors

Intramuscular injections of AAV6 vectors were administered to either 2-day or 42-day old C57BL/6J male mice maintained in a 12:12 h light:dark cycle at 22°C with ad libitum access to normal chow diet and water. Mice were placed under general anesthesia (2% isoflurane in O_2_) and maintained on 37°C heat pads. For injections into 2-day mice, the left hindlimb was injected with rAAV6:MCS (1×10^10^ vector genomes per 10 μl) as a control, and the right contralateral limb was administered rAAV6:LGALS1 (1×10^10^ vector genomes per 10 μl). For injections in *tibialis anterior* (TA) muscle of 42-day old mice, the left muscle was injected with rAAV6:MCS (1×10^9^ to 1×10^10^ vector genomes per 30 μL) as a control, and the right contralateral limb was administered rAAV6:LGALS1 (1×10^9^ to 1×10^10^ vector genomes per 30 μl). At experimental endpoint, mice were humanely killed by cervical dislocation and tissues rapidly excised. Ethical approval for mouse procedures was obtained from The University of Queensland (AEC approval #SBMS/101/13/NHMRC or #SBMS/AIBN/138/16/NHMRC/NHF), QIMR Berghofer Medical Research Institute (AEC approval #A18603M) and the University of Melbourne (AEC approval #1914940).

### Experimental Design and Statistical Rationale

All presented data shows a minimum of three independent biological replicates. For proteomic, glycoproteomic and glycomic analysis of L6 cells, three biological replicates were performed. For glycomic analysis of skeletal muscle, three individual mice per timepoint were analyzed. ANOVA with permutation-based FDR set at 5% with Tukey’s posthoc test was used to determine significantly regulated proteins, glycans or glycopeptides. For generation of clonal CRISPR cells, four independent WT and KO clones were analyzed, and three cover slips analyzed and compared with two-ANOVA. For in vivo AAV delivery, five individual mice were analyzed and compared with paired t-test.

## RESULTS

### Dynamic remodeling of the membrane-associated proteome during *in vitro* myogenesis

To identify membrane-associated proteins potentially important for myogenesis, we performed a quantitative proteomic analysis of L6 myoblasts during differentiation into multinucleated myotubes in 2D cell culture. Proteins were extracted daily over a 7-day time-course of differentiation and membrane-associated proteins were enriched with ultracentrifugation. Proteins were digested with trypsin followed by multiplexed stable isotope labelling with tandem mass tags (TMT) and analysis by 2D liquid chromatography coupled to tandem mass spectrometry (LC-MS/MS) (n=3 biological replicates). A total of 5,898 proteins groups were quantified with at least two peptides represented by 5,801 unique genes in all biological replicates and time-points (**Supplementary Table S1**). More than half of the proteins contained membrane Gene Ontology cellular compartment annotation and broad coverage of proteins associated within the Golgi apparatus and endoplasmic reticulum were also quantified (**Figure 1A**). Principal component analysis revealed clustering of the biological replicates from each of the time points with a rapid and dramatic redistribution of the membrane-associated proteome during differentiation (**Figure 1B**). Remarkably, more than 2,700 proteins were differentially regulated in 7-day myotubes compared to undifferentiated myoblasts (+/− 1.5-fold and q < 0.05 ANOVA with Tukey’s post-hoc test) (**Figure 1C**). Induction of well characterized myogenic markers showed distinct profiles. For example, we observed rapid up-regulation of various troponins (TNNI1/2, TNNC1 and TNNT3) within 1-2 days whereas other proteins associated with the transcriptional regulation such as cysteine and glycine-rich protein-3 (CSRP3) and proteins associated with the contractile apparatus such as dystrophin (DMD) and myosin binding proteins (MYOM1 and MYBPC1) progressively increased throughout the entire differentiation process (**Figure 1D**). KEGG pathway enrichment revealed the regulation of several pathways associated with amino acid, fatty acid and glucose metabolism in addition to other important signaling pathways and proteins associated with *N*-glycosylation (**Figure 1E**). We next performed a clustering analysis of various anabolic and catabolic enzymes associated with glycosylation (**Figure 1F**). We observed up-regulation of various N-acetyl-β-hexosaminidases (HEXA/B/D) and bifunctional UDP-N-acetylglucosamine transferases (ALG13/14) along with the rapid increase of β-galactoside α-2,6-sialyltransferase-1 (ST6GAL1). Conversely, we observed decreased expression of mannosyl glycoprotein acetylglucosaminyl-transferases (MGAT1/2), five isoforms of β-galactoside α-2,3- sialyltransferases (ST3GAL1-5) and various galactosyltransferases (B3GALT6 and B4GALT1/4/5). Taken together, these data will be a rich resource for the study of membrane-associated proteins involved in myogenesis and suggest that the glycosylation machinery, and by extension protein glycosylation, is dynamically regulated.

**Figure 1.**
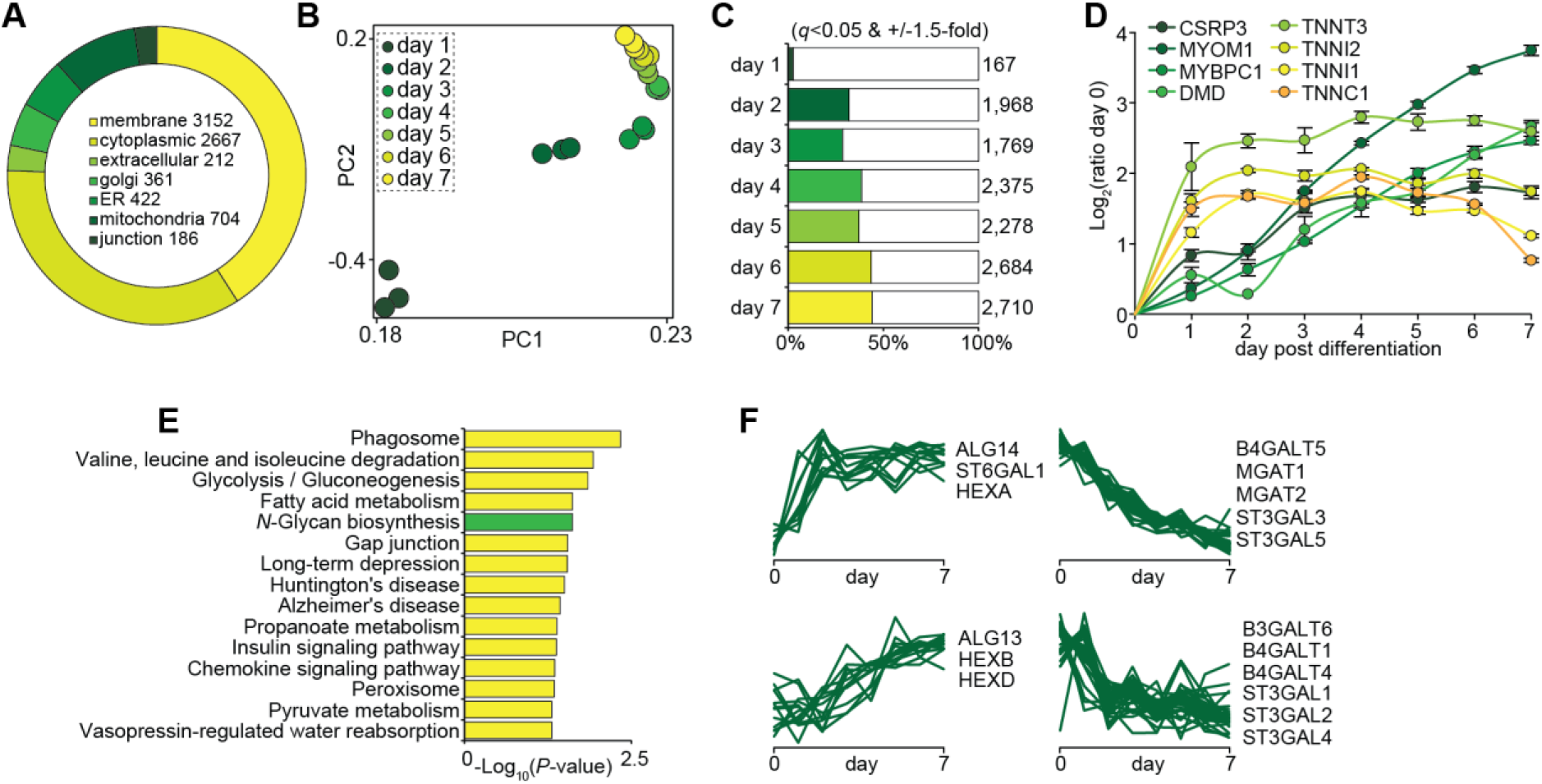
Dynamic membrane-associated proteomic analysis of myogenesis during L6 myotube formation. (**A**) Gene Ontology cellular compartments analysis. (**B**) Principal component analysis. (**C**) Number of regulated protein (ANOVA with permutation-based FDR set at 5% with Tukey’s posthoc test). (**D**) Quantification of myogenic markers. (**E**) KEGG pathway enrichment analysis. (**F**) Temporal clustering of glycosylation-associated proteins.

### Integrated and dynamic *N*-glycomic and *N*-glycoproteomic analysis identifies changes in di-galactosylation to α-2,3-sialylation on cell adhesion and ECM proteins during myogenesis

We next investigated the regulation of protein glycosylation by first performing a glycomic analysis of released *N*-glycans with porous graphitized carbon (PGC) LC-MS/MS over the 7-day time-course of L6 myoblast differentiation (n=3 biological replicates). We identified 45 unique monosaccharide compositions that were confirmed by MS/MS and quantified in all biological replicates and time-points corresponding to a total of 68 different glycan structures (**Supplementary Table S2**). The *N*-glycome was made up of paucimannosidic- (Man1-_3_GlcNAc_2_Fuc0-1), oligomannosidic- (Man_5-9_GlcNAc_2_), hybrid-type glycans capped by a single α- 2,3- or α-2,6-NeuAc or di-Gal moiety with or without core fucosylation, and finally complex-type glycans with mono-, di- or tri-antennae with one to three terminal α-2,3-NeuAc, α-2,6-NeuAc or di-Gal with or without core fucosylation and minor amounts of bisecting β-1,4-GlcNAc (**Supplementary Figure S1**). Analysis of glycans treated with broad-specific sialidase only, a combination of broad-specific sialidase and β-1,4-galactosidase or sialidase and α-galactosidase revealed that the di-Gal comprised a galactose residue α-linked to β-1,4-galactose (**Supplementary Figure S2**). Importantly, none of the di-Gal moieties were found to be capped by sialic acids suggesting that the α-linked Gal is not a substrate for sialyltransfeases in these rat derived cells. A total of 34 glycans were differentially regulated during differentiation (q < 0.05 ANOVA with Tukey’s post-hoc test) (**Figure 2A**). There was progressive decrease in *N*-glycans containing α-2,3-NeuAc (**Figure 2B**) along with a rapid increase in *N*-glycans containing α-2,6- NeuAc (**Figure 2C**). The temporal glycan expression matched the expression of sialyltransferases catalyzing these glycoforms confirming that myoblast-to-myotube differentiation is associated with sialic acid linkage switching (**Figure 2D**). We also observed a progressive decrease in *N*-glycans containing terminal di-Gal epitopes (**Figure 2E**) with a concomitant decrease in the expression of galactosyltransferases during differentiation (**Figure 2F**). Finally, paucimannosylation rapidly increased during differentiation along with the expression of the catalyzing enzyme *N*-acetyl-β-hexosaminidase-A (HEXA) while HEXB only subtly increased (**Figure 2G**).

**Figure 2.**
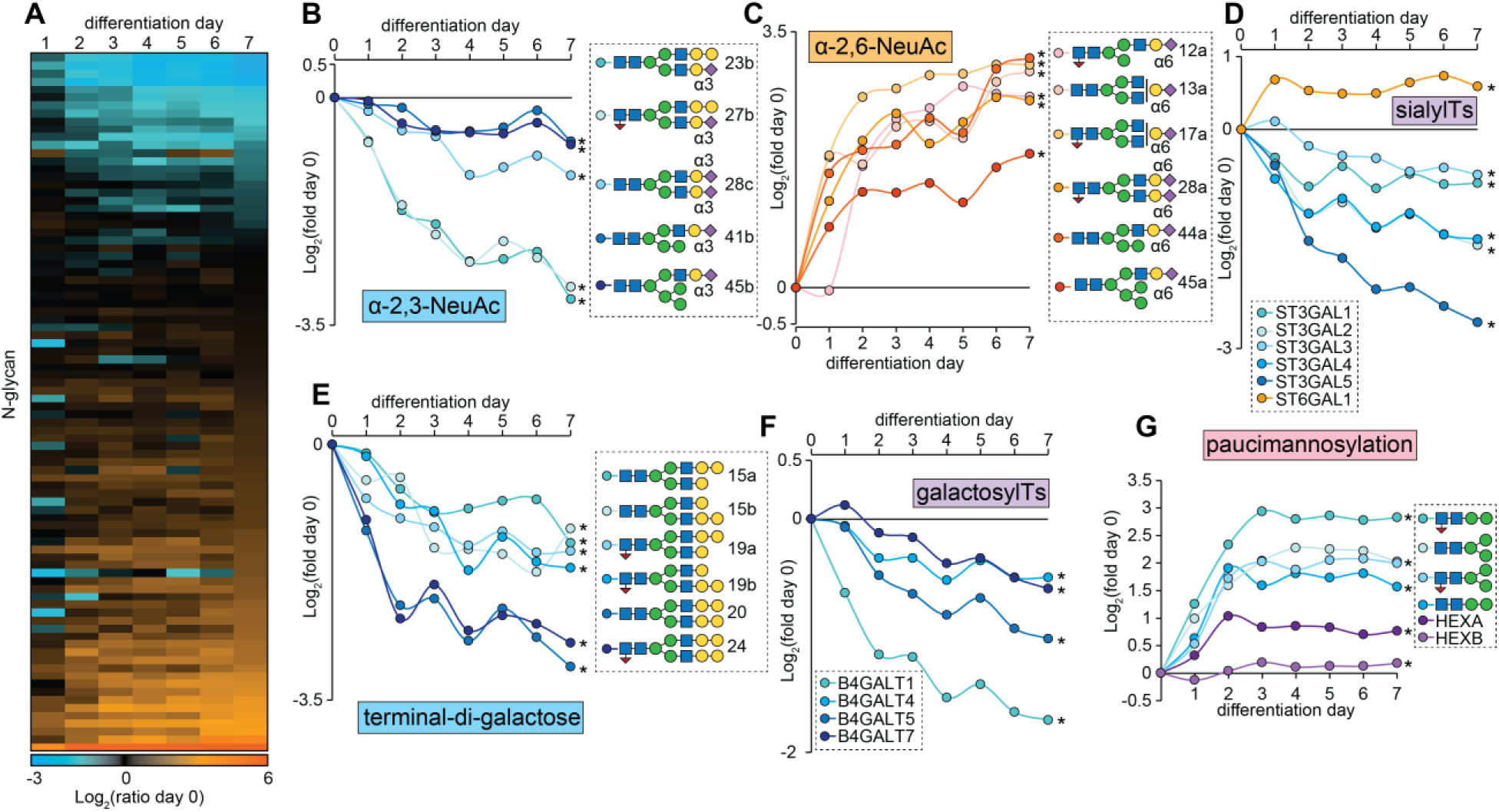
Dynamic *N*-glycomic analysis integrated with proteomic quantification of glycosylation-associated enzymes during L6 myotube formation. (**A**) Heatmap showing the average fold-change of *N*-glycans relative to pre-differentiation at day 0. (**B**) Quantification of α- 2,3-NeuAc-containing *N*-glycans. (**C**) Quantification of α-2,6-NeuAc-containing *N*-glycans. (D) Quantification of sialyltransferases. (E) Quantification of *N*-glycans containing terminal di-Gal. (F) Quantification of galactosyltranferases. (G) Quantification of paucimannosidic *N*-glycans and *N*-acetyl-β-hexosaminidase A/B (HEXA/B). *q < 0.05 ANOVA with permutation-based FDR.

To identify the glycoproteins modified during differentiation we next performed quantitative *N*-glycoproteomics. Membrane-associated proteins were isolated over a 7-day time-course of differentiation (n=3) and digested with trypsin followed by multiplexed stable isotope labelling with TMT. Glycopeptides were enriched with zwitterionic-hydrophilic interaction liquid chromatography (zic-HILIC) and analyzed by LC-MS/MS employing data-dependent fragmentation decisions of higher collisional dissociation (HCD) and product dependent oxonium-ion triggered electron transfer dissociation with HCD supplemental activation (pdEThcD). A total of 2,751 unique glycopeptides were identified with 2,238 identified by HCD and 1,518 identified by pdEThcD (1,005 with both fragmentation types) (**Supplementary Table S3**). A change in the abundance of a specific glycopeptide can arise from either alteration in the: i) site occupancy, ii) type of glycosylation at a given site or, iii) a change in the expression of the glycoprotein itself. To visualize this, **Figure 3A** and **3B** display changes in the abundance of glycopeptides plotted against changes their total protein levels at day 1 and day 7 of differentiation relative to day 0, respectively. The green box highlights glycopeptides from integrin α-3 (ITGA3) which reveals increased protein expression on the y-axis and a distribution of glycoforms on the x-axis. The annotated HCD-MS/MS spectrum indicated with the arrow is shown in **Figure 3C** identifying a glycopeptide containing Asn-605 in the *N*-linked glycosylation motif (NxS/T/C) conjugated with HexNAc(4)Hex(6)Fuc(1) corresponding to a glycan composition with bi-antennary terminal di-Gal moieties. These data are consistent with the global glycomic data showing down regulation of di-Gal-containing *N*-glycans and the observed down regulation of galactosyltransferases in the proteomics data. **Figure 3D** plots the relative changes of this glycopeptide and the total protein levels of ITGA3 across the differentiation time course. ITGA3 was more than 6-fold up-regulated after 7 days of differentiation at the protein level whereas the indicated glycopeptide was almost 2-fold down-regulated. Normalizing these data revealed this specific glycosylation was reduced by more than 10-fold. A total of 931 unique glycopeptides were quantified in at least two biological replicates in all time points and 91 were regulated across the differentiation time course (q < 0.05 ANOVA with Tukey’s post-hoc test) (**Supplementary Table S4**). Several additional cell surface and extracellular proteins contained decreased di-Gal-containing *N*-glycans such as other integrins (ITGA7 and ITGB1), receptors (CD63, ATRN, NOMO1, IL6ST) and extracellular matrix associated proteins (FN1 and EMILIN1). Unlike the glycomic analysis of released *N*-glycans, our analysis of intact glycopeptides is not able to accurately differentiate glycopeptides containing different sialic acid linkages (α-2,3-NeuAc versus α-2,6-NeuAc). Despite this, we observed the regulation of several glycopeptides modified with sialic acid-containing *N*-glycans suggesting changes in the amount of sialylation. These include several proteins also containing of di-galactosylation but additional proteins such as insulin-like growth factor-2 receptor (IGF2R). Many of the proteins with changes in glycosylation including integrins (37) and IGF2R (38) have been previously implicated in the regulation of myogenesis and here we provide support for and expand on these findings by showing precise regulation of di-Gal and sialylation which may play important roles in either protein function or interaction with lectins during the various stage of myogenesis including differentiation, elongation, migration, cell adhesion, membrane alignment or cell fusion.

**Figure 3.**
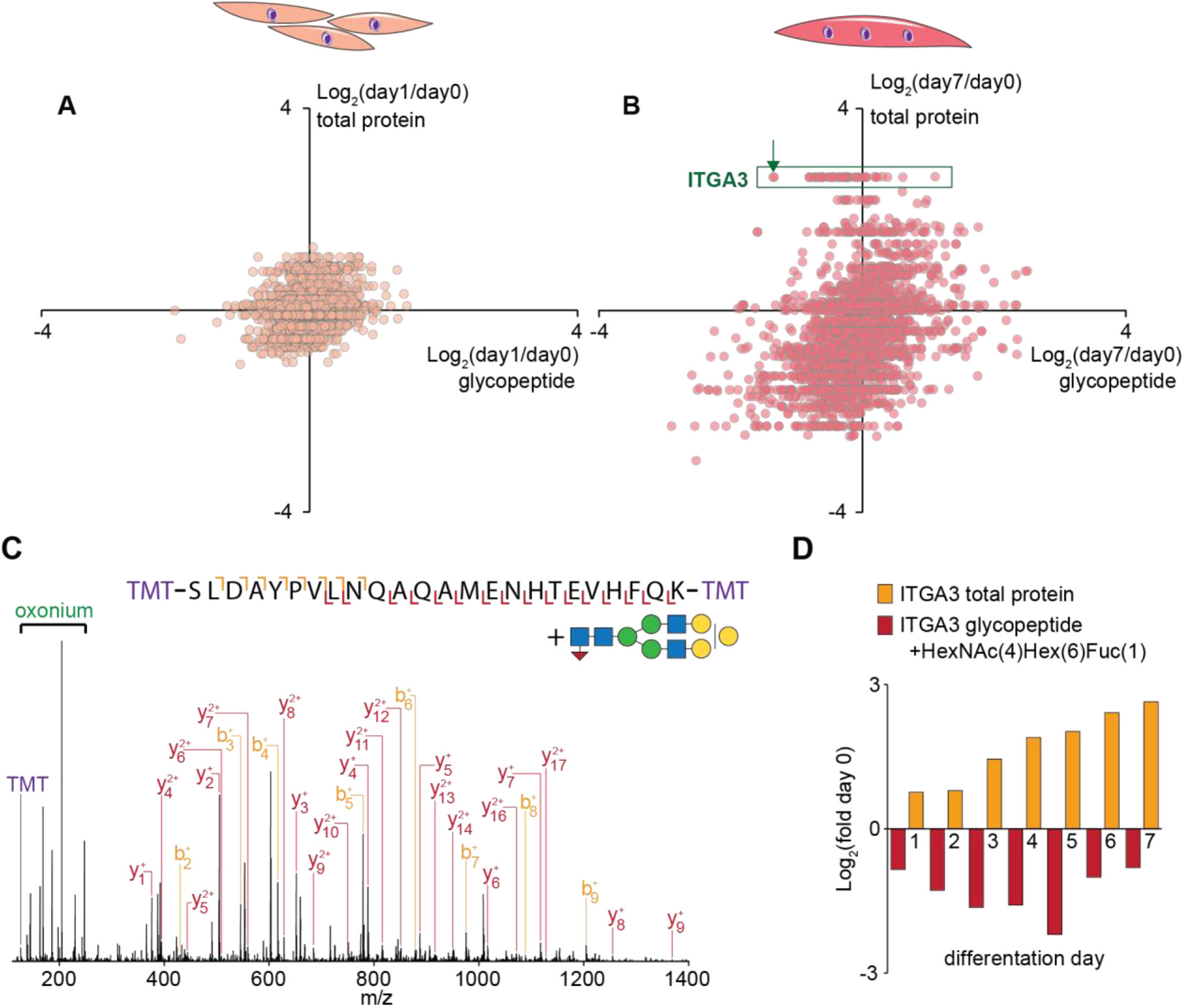
Dynamic *N*-glycoproteomic analysis integrated with proteomic quantification during L6 myotube formation. Quantification of glycopeptides plotted against changes their total protein levels at day 1 (**A**) and day 7 (**B**) of differentiation relative to day 0. The green box highlights glycopeptides from integrin α-3 (ITGA3). Arrow indicates annotated HCD-MS/MS spectrum in (**C**) identifying a glycopeptide containing Asn605 Win the *N*-glycosylation motif (NxS/T/C) with the glycan HexNAc(4)Hex(6)Fuc(1) corresponding to a glycan composition with bi-antennary terminal di-Gal. (**D**) Relative changes of the indicated glycopeptide and the total protein levels of ITGA3 across the differentiation time-course.

### Galectin-1 is required for myoblast differentiation *in vitro*

Many glycoproteins form glycan:protein interactions with GBPs (39). The regulation of galactosylation during myogenesis is interesting as these glycans can be recognized by various galectins, forming interactions that play important roles in several biological processes (40). Mice lacking various galectins display alterations in smooth muscle, cardiac muscle and skeletal muscle development or regeneration/remodeling following injury (10, 41, 42). However, galectins are ubiquitously expressed in many cell types and their roles in the various stages of myogenesis and muscle development are incompletely understood. Our proteomic analysis identified time-dependent regulation of several galectins during the differentiation of L6 myoblasts *in vitro* including the up-regulation of galectin-1 (LGALS1) and down-regulation of galectin-3 (LGALS3) (**Figure 4A**). We also observed an initial subtle increase in galectin-8 (LGALS8) which declined after 2 days while galectin-3 binding protein (LGALS3BP) initially decreased and then increased after 3 days of differentiation. Here, we focus on galectin-1 and confirm time-dependent increases in protein expression during differentiation of L6 myoblasts by western blot analysis (**Figure 4B**). We next genetically ablated *Lgals1* from L6 myoblasts using paired gRNAs and CRISPR/Cas9(D10A) (**Figure 4C**). Clonal cells were isolated and four independent knockout (KO) lines were validated by western-blot analysis (**Figure 4D)**. We also isolated four wildtype (WT) clonal cells from the original population transfected with the gRNAs and CRISPR/Cas9(D10A) that did not undergo editing to serve as controls. After 3 days of differentiation, WT myoblasts formed elongated multinucleated and myosin positive myotubes while KO cells mostly remained mononucleated with a small number of multinucleated cells which stained for myosin (**Figure 4E**). Quantification of the differentiation index showed a marked decrease in the percentage of nuclei in myosin positive cells indicating loss of *Lgals1* significantly reduced myoblast differentiation (**Figure 4F**). Our results support previous *in vivo* data showing mice lacking *Lgals1* have reduced myofiber diameter and delayed muscle regeneration (10), and we propose impaired myotube formation is likely mediated by an early defect in the myogenic program and failure to initiate differentiation.

**Figure 4.**
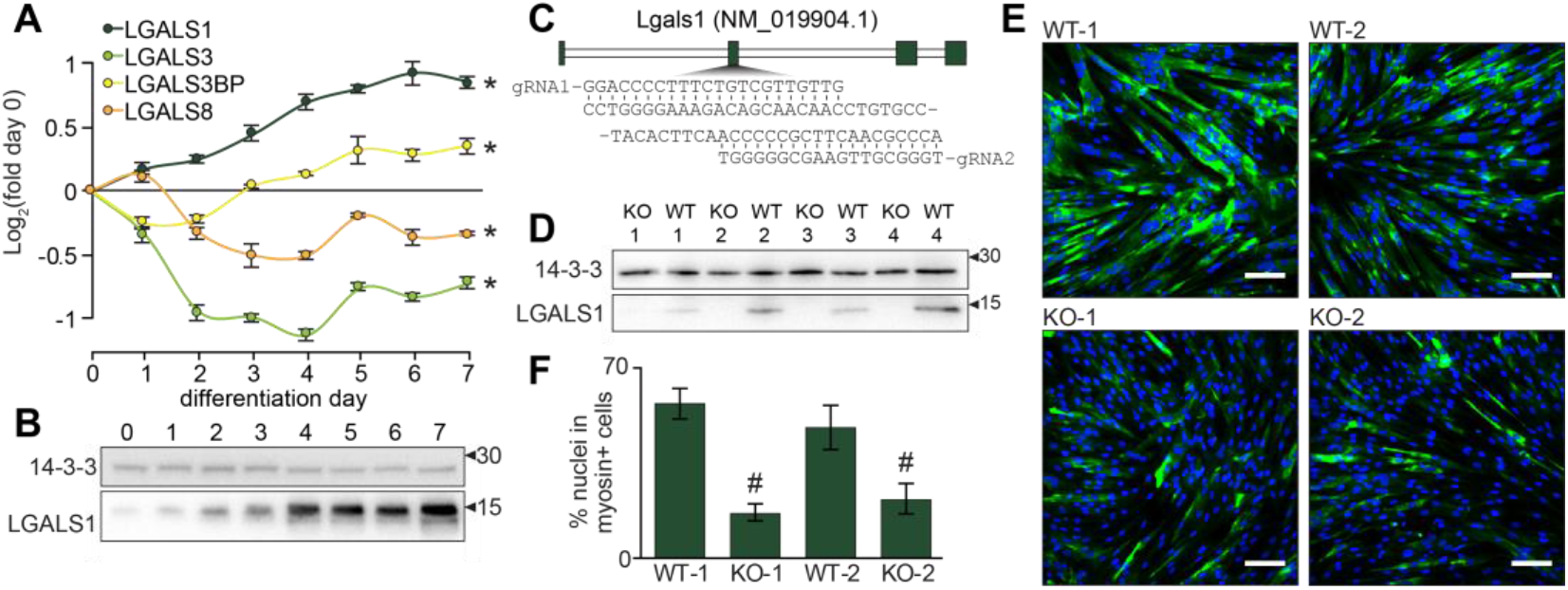
Galectin-1 is up-regulated and required for L6 myotube formation. (**A**) Quantification of galectins during myogenesis. (**B**) Western-blot analysis of galectin-1 during myogenesis. (**C**) CRISPR/Cas9(D10A) paired gRNA strategy to knockout *Lgals1*. (**D**) Western-blot analysis of galectin-1 and 14-3-3 loading control in wild-type (WT) and knockout (KO) L6 myoblasts. (**E**) Immunofluorescence microscopy of WT and KO cells at day 3 of differentiation (blue = Hoechst; green = myosin; white scale bar = 50 μm). (**F**) Quantification of the differentiation index expressed as the percentage of nuclei in myosin positive cells. *q < 0.05 ANOVA with permutation-based FDR. #p < 0.05 two-way ANOVA.

### Galectin-1 and *N*-glycosylation are regulated during mouse muscle development *in vivo*

We next investigated the regulation of protein *N*-glycosylation and galectin-1 expression during postnatal mouse skeletal muscle development. Male mice were sacrificed at 1, 7, 14 and 21 days after birth and skeletal muscle collected from lower hindlimbs (n=3 biological replicates). Western-blot analysis revealed an increase in galectin-1 throughout development consistent with the *in vitro* myogenesis data (**Figure 5A**). Next, released *N*-glycans were quantified in total protein lysates throughout postnatal development with PGC-LC-MS/MS. We identified 54 unique monosaccharide compositions that were confirmed by MS/MS and quantified in all biological replicates and developmental time-points corresponding to a total of 104 different glycan structures (**Supplementary Table S5**). The *N*-glycome was made up of paucimannosidic-, oligomannosidic-, and hybrid-type glycans capped with a single α-2,3- or α-2,6-NeuAc or di-Gal moiety with or without core fucosylation, and also complex-type glycans displaying mono-, di-or tri-antennae with one or two terminal α-2,3-NeuAc, α-2,6-NeuAc, α-2,3-NeuGc, α-2,6-NeuGc or di-Gal with or without core fucosylation (**Supplementary Figure S3**). There was a progressive decrease in N-glycans containing α-2,3-NeuAc that was significant 21 days after birth (**Figure 5C**). Furthermore, a rapid reduction in α-2,6-NeuAc was also observed 7 days after birth but this gradually increased throughout the next two weeks of development. The seemingly inverse relationship between these two sialic acid linkages is consistent with the *in vitro* myogenesis model and the biological consequences of these changes on the myogenic program warrants further investigation. Also in line with the *in vitro* data, the *N*-glycans containing terminal di-Gal were reduced at day 7, 14 and 21 days after birth and paucimannosylation was significantly increased 14 days after birth and by 21 days the levels were >2.5-fold greater than new-born mice. Taken together, our data suggest that *N*-glycan remodeling is a strong feature associated with *in vitro* and *in vivo* rodent models of myogenesis.

**Figure 5.**
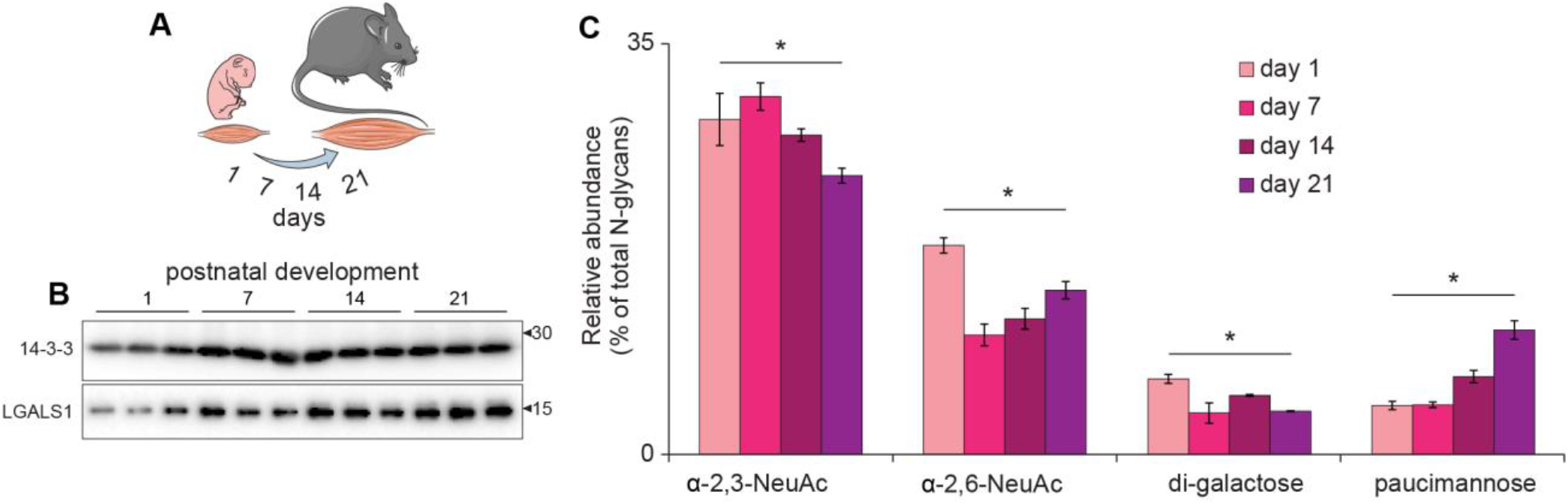
Quantification of LGALS1 and released *N*-glycans during postnatal skeletal muscle development in mice. (**A**) Postnatal time-points. (**B**) Western blot analysis of galecin-1 and 14-3-3 loading control in skeletal muscle development. (**C**) Quantification of selected *N*- glycan features during skeletal muscle development. *q < 0.05 ANOVA with permutation-based FDR.

### *In vivo* galectin-1 gain-of-function increases muscle mass

Administration of recombinant galectin-1 has recently been shown to improve muscle function in a mouse model of muscular dystrophy (14). Therefore, we next investigated the ability of galectin-1 to promote postnatal muscle development using an *in vivo* gene therapy gain-of-function mouse model. Hindlimbs of new-born mice were injected with recombinant adeno-associated virus serotype-6 (rAAV6) employing a paired experimental design where the left legs were treated with viral vectors encoding an empty multiple cloning site (MCS) to severe as a control and the right legs received constructs expressing *Lgals1* to transduce the entire lower limb musculature (n=5 biological replicates) (**Figure 6A**). An initial pilot dose response was performed in the *tibialis anterior* (TA) muscle of 42-day old mice revealing injections of ~1 x 10^10^ vector genomes (vg) produced ~3-fold average overexpression by western-blot analysis (data not shown). Two days after birth, mice were injected with rAAV6 and following 42-days the animals were sacrificed and muscles dissected. Western-blot analysis of *gastrocnemius* muscle revealed a variable but significant ~2.2-fold average overexpression of galectin-1 (**Figure 6B-C**). The TA muscles injected with rAAV6:LGALS1 all increased in mass with an average of 5.3% but there was considerable variation likely due to variation in expression (**Figure 6D**). Our data is the first to reveal that enhanced muscle is mass associated with galecin-1 expression in early skeletal muscle development.

**Figure 6.**
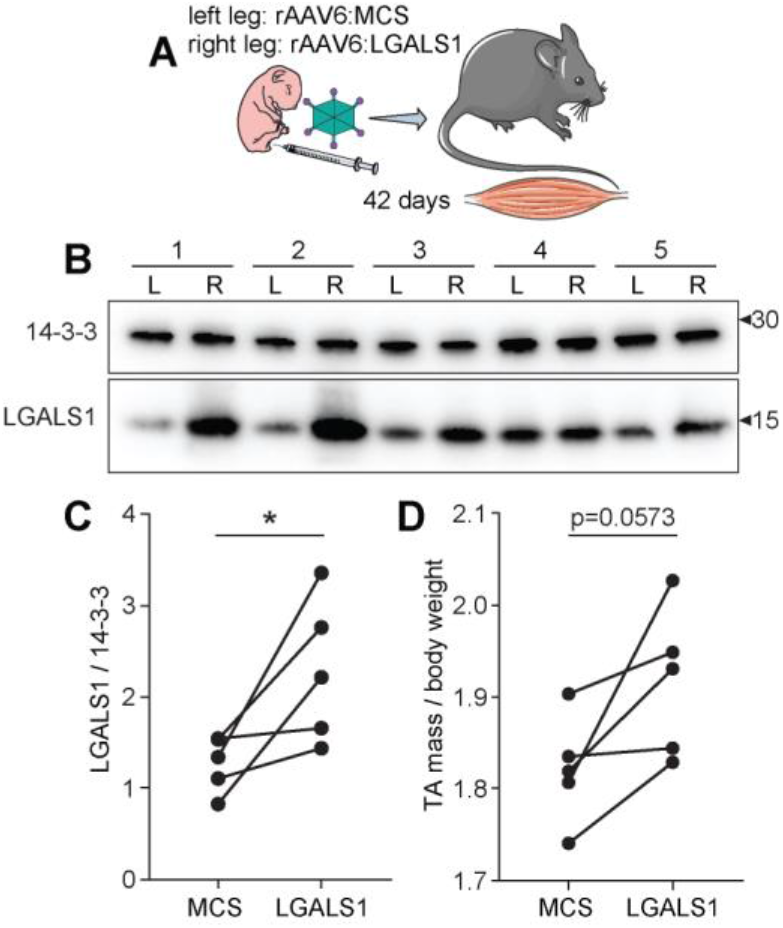
*In vivo* galectin-1 overexpression increases postnatal muscle mass. (A) Strategy for *in vivo* gain-of-function mouse model using recombinant adeno-associated virus serotype-6 (rAAV6) to express an empty multiple cloning site (MCS) control or LGALS1. (B) Western blot analysis of galectin-1 and 14-3-3 loading control of skeletal muscle from left leg (L) injected with rAAV6:MCS control and right leg (R) injected with rAAV6:LGALS1. (C) Quantification of western blot. (D) Mass of *tibialis anterior* (TA) muscle normalized to body weight injected with rAAV6:MCS control or rAAV6:LGALS1. *p < 0.05 paired t-test.

## DISCUSSION

The integration of proteomics, glycoproteomics and glycomics is proving to be a valuable approach for holistic characterization of glycoproteome regulation (43). No single technology can comprehensively unravel the complexities of glycosylation and each platform offers unique insights; proteomics enables quantification of glycoproteins, lectins and glycosyltransferase/glycosidase abundance; glycoproteomics enables quantification of site-specific changes in glycan composition; and glycomics enables detailed structural characterization of the glycome. Here, we integrate these technologies to study dynamic regulation of protein *N*-glycosylation enabling us to define changes in the abundance of galectins, galactosyltransferases, sialyltransferases and *N*-acetyl-β-hexosaminidases that correlate with specific glycan structures on functionally-important receptors and adhesion molecules. These integrated data substantially improve our confidence in defining the mechanisms of glycosylation changes during myogenesis. Specifically, we show a switch in terminal di-galactosylation and sialylation linkages mediated by changes in glycosyltransferase/sialyltransferase expression. The is interesting because these enzymes are co-localized in the Golgi Apparatus and will compete for the generation of galacto-or sialo-epitopes presented for reactivity. It is important to note that di-galactosylation is not a common feature in the human glycome meaning the use of these glycoepitopes in myogenesis may be different to rodent models analyzed in the current study. Our data also revealed an increase in paucimannosylation during terminal differentiation which is surprising as this modification has previously been associated with proliferation in cancer (44) and neuronal progenitor cells (45), and is down-regulated during macrophage differentiation (46).

Recent advances in glycoproteomics are dramatically increasing our ability to identify thousands of glycopeptides (47–54), and future studies are likely to see an expansion of methods to analyze large sample cohorts and further enhance identification confidence or structural characterization (55–57). One missing piece of information in our study is the assessment of glycan heterogeneity and site-specific occupancy on intact protein species. New methods focused on top-down glycoproteomics to characterize intact glycoproteins will further enhance our ability to achieve this level of detail on a proteome-wide scale (58). Our study focused on *N*-glycosylation but it is important to note that other types of protein glycosylation such as *O*-, *GPI*- or *C*-linked may also be important during myogenesis. Furthermore, additional glycoconjugates such as glycosphingolipids, glycosaminoglycans and proteoglycans are likely to play important roles in the development of skeletal muscle as mutations in the catalytic enzymes responsible for their biosynthesis cause developmental neurological defects (59), and pharmacological inhibition modulates adult skeletal muscle metabolic function (60).

Our proteomic analysis quantified an increase in galecin-1 abundance and a reciprocal down-regulation of galectin-3. This is interesting because both galectins have similar functions such as the regulation of Ras-mediated signaling in cancer (61). Reciprocal regulation of galectin-1 and galectin-3 has also been observed in sera from rheumatoid arthritis patients undergoing various treatments (62) suggesting possible compensatory mechanisms. Galectins form a family of 15 members that bind to galactose via conserved carbohydrate binding domains (CBDs) (63). Galectin-1 is a prototypical galectin containing a single CBD that form dimers while galectin-3 contains an extended N-terminal domain with the ability to form pentamers (64). Administration of recombinant galectin-1 to the mdx mouse model of muscle dystrophy increased utrophin and integrin expression, and improved skeletal muscle function (14). Furthermore, enhanced muscle function has also been observed with exogenous galectin-3 expression in a similar mdx mouse model (65). Our data further support these observations revealing overexpression of galectin-1 increases muscle mass during postnatal muscle development in mice. Galectins have been shown to bind galecto-epitopes on various cell surface receptors including integrins to promote signaling (63). Surprisingly, we show that various integrins display decreased di-galactosylation during myogenesis suggesting reduced galectin binding ability. This is somewhat contradictory however, the presence of α-2,3-NeuAc vs α-2,6-NeuAc can further influence the binding of various galectins to galactose epitopes. For example, α-2,6-NeuAc inhibits galectin-1 binding to galactose but has no effect on galectin-3 binding while both galectin-1 and −3 can bind galactose containing α-2,3-NeuAc-linked glycans (66). The differential binding of various galectins to glycans containing different sialic acid linkage isomers further supports a complex interplay between these glycoforms in the various stages of myogenesis. Further experiments are required to precisely quantify the glycan-protein interactions during myogenesis and muscle growth. Galectins are secreted via non-classical secretory mechanisms and may also play a role in endocytosis (67, 68). Therefore, galectins have been primarily studied in the context of extracellular lattice formation however, there is growing evidence to suggest galectins may regulate intracellular signaling via lysosomal trafficking (69). For example, LGALS8 has recently been shown to inhibit mTOR signaling at the lysosome (70). This is interesting because the activation of mTOR has well established roles in skeletal muscle growth (71). Our data revealed a decrease in LGALS8 expression during myogenesis suggesting a relief in mTOR inhibition required for cell growth and further experiments are required to investigate this in skeletal muscle. We anticipate our integrated analysis of proteomics, glycomics and glycoproteomics will be a valuable resource to further our understanding of the role of glycosylation on muscle development.

## Abbreviations

AAV: adeno-associated virus
ANOVA: analysis of variation
CBDs: carbohydrate binding domains
CDGs: congenital disorders of glycosylation
CRISPR: clustered regularly interspaced short palindromic repeats
CID: collisional induced dissociation
DTT: dithiothreitol
ECM: extracellular matrix
EThcD: electron transfer dissociation with higher collisional dissociation supplemental activation
FBS: fetal bovine serum
FDR: false discovery rate
FA: formic acid
GBPs: glycan-binding proteins
HCD: higher collisional dissociation
HILIC: hydrophilic interaction liquid chromatography
MCS: multiple cloning site
MeCN: acetonitrile
NMJ: neuromuscular junctions
PCA: principal component analysis
PGC: porous graphitized carbon
PSM: peptide spectral match
TEAB: triethylammonium bicarbonate
TFA: trifluoroacetic acid
TMT: tandem mass tags

## ACKNOWLEDGEMENTS

We thank David James and the members of the Metabolic Systems Biology group for feedback and discussions. This research was facilitated by access to Sydney Mass Spectrometry, a core research facility at the University of Sydney. This work was supported by infrastructure and technical assistance from the Melbourne Mouse Metabolic Phenotyping Platform (MMMPP) at the University of Melbourne. This work was funded by an NHMRC Early Career Fellowship and the University of Melbourne Driving Research Momentum program (B.L.P). J.E.H. is supported by grants and fellowships from NHMRC and National Heart Foundation of Australia. The contents of the published material are solely the responsibility of the individual authors and do not reflect the view of NHMRC.

## DATA AVAILABILITY

Supplementary data is available online. Proteomic and glycoproteomic data are available via ProteomeXchange with identifier PXD019372 (23). Username: and Password: KGAgOAkh. Glycomic data are freely available on Glycopost (32) with the identifier GPST000079 (raw files fhttps://glycopost.glycosmos.org/preview/7030709905ed7e73ea0928, PIN: 5219) and Panorama Public (33) (skyline assays, https://panoramaweb.org/RatMuscleGlyco.url).

## REFERENCES

1. Dittmar, T., and Zanker, K. S. (2011) Cell fusion in health and disease. Volume II: cell fusion in disease. Introduction. Adv Exp Med Biol 714, 1–3

2. Braun, T., and Gautel, M. (2011) Transcriptional mechanisms regulating skeletal muscle differentiation, growth and homeostasis. Nat Rev Mol Cell Biol 12, 349–361

3. Kim, J. H., Jin, P., Duan, R., and Chen, E. H. (2015) Mechanisms of myoblast fusion during muscle development. Curr Opin Genet Dev 32, 162–170

4. Kablar, B., and Rudnicki, M. A. (2000) Skeletal muscle development in the mouse embryo. Histol Histopathol 15, 649–656

5. Schiaffino, S., Dyar, K. A., Ciciliot, S., Blaauw, B., and Sandri, M. (2013) Mechanisms regulating skeletal muscle growth and atrophy. FEBS J 280, 4294–4314

6. Scott, K., Gadomski, T., Kozicz, T., and Morava, E. (2014) Congenital disorders of glycosylation: new defects and still counting. J Inherit Metab Dis 37, 609–617

7. Grunewald, S. (2009) The clinical spectrum of phosphomannomutase 2 deficiency (CDG-Ia). Biochim Biophys Acta 1792, 827–834

8. Shrimal, S., Ng, B. G., Losfeld, M. E., Gilmore, R., and Freeze, H. H. (2013) Mutations in STT3A and STT3B cause two congenital disorders of glycosylation. Hum Mol Genet 22, 4638–4645

9. Rymen, D., Peanne, R., Millon, M. B., Race, V., Sturiale, L., Garozzo, D., Mills, P., Clayton, P., Asteggiano, C. G., Quelhas, D., Cansu, A., Martins, E., Nassogne, M. C., Goncalves-Rocha, M., Topaloglu, H., Jaeken, J., Foulquier, F., and Matthijs, G. (2013) MAN1B1 deficiency: an unexpected CDG-II. PLoS Genet 9, e1003989

10. Georgiadis, V., Stewart, H. J., Pollard, H. J., Tavsanoglu, Y., Prasad, R., Horwood, J., Deltour, L., Goldring, K., Poirier, F., and Lawrence-Watt, D. J. (2007) Lack of galectin-1 results in defects in myoblast fusion and muscle regeneration. Dev Dyn 236, 1014–1024

11. Ahmed, H., Du, S. J., and Vasta, G. R. (2009) Knockdown of a galectin-1-like protein in zebrafish (Danio rerio) causes defects in skeletal muscle development. Glycoconj J 26, 277–283

12. Gu, M., Wang, W., Song, W. K., Cooper, D. N., and Kaufman, S. J. (1994) Selective modulation of the interaction of alpha 7 beta 1 integrin with fibronectin and laminin by L-14 lectin during skeletal muscle differentiation. J Cell Sci 107 (Pt 1), 175–181

13. McMorran, B. J., Miceli, M. C., and Baum, L. G. (2017) Lectin-binding characterizes the healthy human skeletal muscle glycophenotype and identifies disease-specific changes in dystrophic muscle. Glycobiology 27, 1134–1143

14. Wuebbles, R. D., Cruz, V., Van Ry, P., Barraza-Flores, P., Brewer, P. D., Jones, P., and Burkin, D. J. (2019) Human Galectin-1 Improves Sarcolemma Stability and Muscle Vascularization in the mdx Mouse Model of Duchenne Muscular Dystrophy. Mol Ther Methods Clin Dev 13, 145–153

15. Parker, B. L., Thaysen-Andersen, M., Fazakerley, D. J., Holliday, M., Packer, N. H., and James, D. E. (2016) Terminal Galactosylation and Sialylation Switching on Membrane Glycoproteins upon TNF-Alpha-Induced Insulin Resistance in Adipocytes. Mol Cell Proteomics 15, 141–153

16. Palmisano, G., Lendal, S. E., Engholm-Keller, K., Leth-Larsen, R., Parker, B. L., and Larsen, M. R. (2010) Selective enrichment of sialic acid-containing glycopeptides using titanium dioxide chromatography with analysis by HILIC and mass spectrometry. Nat Protoc 5, 1974–1982

17. Hagglund, P., Bunkenborg, J., Elortza, F., Jensen, O. N., and Roepstorff, P. (2004) A new strategy for identification of N-glycosylated proteins and unambiguous assignment of their glycosylation sites using HILIC enrichment and partial deglycosylation. J Proteome Res 3, 556–566

18. Mysling, S., Palmisano, G., Hojrup, P., and Thaysen-Andersen, M. (2010) Utilizing ion-pairing hydrophilic interaction chromatography solid phase extraction for efficient glycopeptide enrichment in glycoproteomics. Anal Chem 82, 5598–5609

19. McAlister, G. C., Nusinow, D. P., Jedrychowski, M. P., Wuhr, M., Huttlin, E. L., Erickson, B. K., Rad, R., Haas, W., and Gygi, S. P. (2014) MultiNotch MS3 enables accurate, sensitive, and multiplexed detection of differential expression across cancer cell line proteomes. Anal Chem 86, 7150–7158

20. Frese, C. K., Altelaar, A. F., van den Toorn, H., Nolting, D., Griep-Raming, J., Heck, A. J., and Mohammed, S. (2012) Toward full peptide sequence coverage by dual fragmentation combining electron-transfer and higher-energy collision dissociation tandem mass spectrometry. Anal Chem 84, 9668–9673

21. Saba, J., Dutta, S., Hemenway, E., and Viner, R. (2012) Increasing the productivity of glycopeptides analysis by using higher-energy collision dissociation-accurate mass-product-dependent electron transfer dissociation. Int J Proteomics 2012, 560391

22. Wu, S. W., Pu, T. H., Viner, R., and Khoo, K. H. (2014) Novel LC-MS(2) product dependent parallel data acquisition function and data analysis workflow for sequencing and identification of intact glycopeptides. Anal Chem 86, 5478–5486

23. Perez-Riverol, Y., Csordas, A., Bai, J., Bernal-Llinares, M., Hewapathirana, S., Kundu, D. J., Inuganti, A., Griss, J., Mayer, G., Eisenacher, M., Perez, E., Uszkoreit, J., Pfeuffer, J., Sachsenberg, T., Yilmaz, S., Tiwary, S., Cox, J., Audain, E., Walzer, M., Jarnuczak, A. F., Ternent, T., Brazma, A., and Vizcaino, J. A. (2019) The PRIDE database and related tools and resources in 2019: improving support for quantification data. Nucleic Acids Res 47, D442–D450

24. MacCoss, M. J., Wu, C. C., and Yates, J. R., 3rd (2002) Probability-based validation of protein identifications using a modified SEQUEST algorithm. Anal Chem 74, 5593–5599

25. Kall, L., Canterbury, J. D., Weston, J., Noble, W. S., and MacCoss, M. J. (2007) Semi-supervised learning for peptide identification from shotgun proteomics datasets. Nat Methods 4, 923–925

26. Johnson, W. E., Li, C., and Rabinovic, A. (2007) Adjusting batch effects in microarray expression data using empirical Bayes methods. Biostatistics 8, 118–127

27. Bern, M., Kil, Y. J., and Becker, C. (2012) Byonic: advanced peptide and protein identification software. Curr Protoc Bioinformatics Chapter 13, Unit13 20

28. Lee, L. Y., Moh, E. S., Parker, B. L., Bern, M., Packer, N. H., and Thaysen-Andersen, M. (2016) Toward Automated N-Glycopeptide Identification in Glycoproteomics. J Proteome Res 15, 3904–3915

29. Parker, B. L., Thaysen-Andersen, M., Solis, N., Scott, N. E., Larsen, M. R., Graham, M. E., Packer, N. H., and Cordwell, S. J. (2013) Site-specific glycan-peptide analysis for determination of N-glycoproteome heterogeneity. J Proteome Res 12, 5791–5800

30. Jensen, P. H., Karlsson, N. G., Kolarich, D., and Packer, N. H. (2012) Structural analysis of N-and O-glycans released from glycoproteins. Nat Protoc 7, 1299–1310

31. Adams, K. J., Pratt, B., Bose, N., Dubois, L. G., St John-Williams, L., Perrott, K. M., Ky, K., Kapahi, P., Sharma, V., MacCoss, M. J., Moseley, M. A., Colton, C. A., MacLean, B. X., Schilling, B., Thompson, J. W., and Alzheimer’s Disease Metabolomics, C. (2020) Skyline for Small Molecules: A Unifying Software Package for Quantitative Metabolomics. J Proteome Res 19, 1447–1458

32. Varki, A., Cummings, R. D., Aebi, M., Packer, N. H., Seeberger, P. H., Esko, J. D., Stanley, P., Hart, G., Darvill, A., Kinoshita, T., Prestegard, J. J., Schnaar, R. L., Freeze, H. H., Marth, J. D., Bertozzi, C. R., Etzler, M. E., Frank, M., Vliegenthart, J. F., Lutteke, T., Perez, S., Bolton, E., Rudd, P., Paulson, J., Kanehisa, M., Toukach, P., Aoki-Kinoshita, K. F., Dell, A., Narimatsu, H., York, W., Taniguchi, N., and Kornfeld, S. (2015) Symbol Nomenclature for Graphical Representations of Glycans. Glycobiology 25, 1323–1324

33. Rojas-Macias, M. A., Mariethoz, J., Andersson, P., Jin, C., Venkatakrishnan, V., Aoki, N. P., Shinmachi, D., Ashwood, C., Madunic, K., Zhang, T., Miller, R. L., Horlacher, O., Struwe, W. B., Watanabe, Y., Okuda, S., Levander, F., Kolarich, D., Rudd, P. M., Wuhrer, M., Kettner, C., Packer, N. H., Aoki-Kinoshita, K. F., Lisacek, F., and Karlsson, N. G. (2019) Towards a standardized bioinformatics infrastructure for N-and O-glycomics. Nat Commun 10, 3275

34. Sharma, V., Eckels, J., Schilling, B., Ludwig, C., Jaffe, J. D., MacCoss, M. J., and MacLean, B. (2018) Panorama Public: A Public Repository for Quantitative Data Sets Processed in Skyline. Mol Cell Proteomics 17, 1239–1244

35. Schneider, C. A., Rasband, W. S., and Eliceiri, K. W. (2012) NIH Image to ImageJ: 25 years of image analysis. Nat Methods 9, 671–675

36. Blankinship, M. J., Gregorevic, P., Allen, J. M., Harper, S. Q., Harper, H., Halbert, C. L., Miller, A. D., and Chamberlain, J. S. (2004) Efficient transduction of skeletal muscle using vectors based on adeno-associated virus serotype 6. Mol Ther 10, 671–678

37. Boppart, M. D., and Mahmassani, Z. S. (2019) Integrin signaling: linking mechanical stimulation to skeletal muscle hypertrophy. Am J Physiol Cell Physiol 317, C629–C641

38. Torrente, Y., Bella, P., Tripodi, L., Villa, C., and Farini, A. (2020) Role of Insulin-Like Growth Factor Receptor 2 across Muscle Homeostasis: Implications for Treating Muscular Dystrophy. Cells 9

39. (2015) In: rd, Varki, A., Cummings, R. D., Esko, J. D., Stanley, P., Hart, G. W., Aebi, M., Darvill, A. G., Kinoshita, T., Packer, N. H., Prestegard, J. H., Schnaar, R. L., and Seeberger, P. H., eds. Essentials of Glycobiology, Cold Spring Harbor (NY)

40. Di Lella, S., Sundblad, V., Cerliani, J. P., Guardia, C. M., Estrin, D. A., Vasta, G. R., and Rabinovich, G. A. (2011) When galectins recognize glycans: from biochemistry to physiology and back again. Biochemistry 50, 7842–7857

41. Tsai, M. S., Chiang, M. T., Tsai, D. L., Yang, C. W., Hou, H. S., Li, Y. R., Chang, P. C., Lin, H. H., Chen, H. Y., Hwang, I. S., Wei, P. K., Hsu, C. P., Lin, K. I., Liu, F. T., and Chau, L. Y. (2018) Galectin-1 Restricts Vascular Smooth Muscle Cell Motility Via Modulating Adhesion Force and Focal Adhesion Dynamics. Sci Rep 8, 11497

42. Nguyen, M. N., Ziemann, M., Kiriazis, H., Su, Y., Thomas, Z., Lu, Q., Donner, D. G., Zhao, W. B., Rafehi, H., Sadoshima, J., McMullen, J. R., El-Osta, A., and Du, X. J. (2019) Galectin-3 deficiency ameliorates fibrosis and remodeling in dilated cardiomyopathy mice with enhanced Mst1 signaling. Am J Physiol Heart Circ Physiol 316, H45–H60

43. Raghunathan, R., Sethi, M. K., Klein, J. A., and Zaia, J. (2019) Proteomics, Glycomics, and Glycoproteomics of Matrisome Molecules. Mol Cell Proteomics 18, 2138–2148

44. Chatterjee, S., Lee, L. Y., Kawahara, R., Abrahams, J. L., Adamczyk, B., Anugraham, M., Ashwood, C., Sumer-Bayraktar, Z., Briggs, M. T., Chik, J. H. L., Everest-Dass, A., Forster, S., Hinneburg, H., Leite, K. R. M., Loke, I., Moginger, U., Moh, E. S. X., Nakano, M., Recuero, S., Sethi, M. K., Srougi, M., Stavenhagen, K., Venkatakrishnan, V., Wongtrakul-Kish, K., Diestel, S., Hoffmann, P., Karlsson, N. G., Kolarich, D., Molloy, M. P., Muders, M. H., Oehler, M. K., Packer, N. H., Palmisano, G., and Thaysen-Andersen, M. (2019) Protein Paucimannosylation Is an Enriched N-Glycosylation Signature of Human Cancers. Proteomics 19, e1900010

45. Dahmen, A. C., Fergen, M. T., Laurini, C., Schmitz, B., Loke, I., Thaysen-Andersen, M., and Diestel, S. (2015) Paucimannosidic glycoepitopes are functionally involved in proliferation of neural progenitor cells in the subventricular zone. Glycobiology 25, 869–880

46. Hinneburg, H., Pedersen, J. L., Bokil, N. J., Pralow, A., Schirmeister, F., Kawahara, R., Rapp, E., Saunders, B. M., and Thaysen-Andersen, M. (2020) High-resolution longitudinal N-and O-glycoprofiling of human monocyte-to-macrophage transition. Glycobiology

47. Stadlmann, J., Taubenschmid, J., Wenzel, D., Gattinger, A., Durnberger, G., Dusberger, F., Elling, U., Mach, L., Mechtler, K., and Penninger, J. M. (2017) Comparative glycoproteomics of stem cells identifies new players in ricin toxicity. Nature 549, 538–542

48. Riley, N. M., Hebert, A. S., Westphall, M. S., and Coon, J. J. (2019) Capturing site-specific heterogeneity with large-scale N-glycoproteome analysis. Nat Commun 10, 1311

49. Zhang, Y., Mao, Y., Zhao, W., Su, T., Zhong, Y., Fu, L., Zhu, J., Cheng, J., and Yang, H. (2020) Glyco-CPLL: An Integrated Method for In-Depth and Comprehensive N-Glycoproteome Profiling of Human Plasma. J Proteome Res 19, 655–666

50. Shu, Q., Li, M., Shu, L., An, Z., Wang, J., Lv, H., Yang, M., Cai, T., Hu, T., Fu, Y., and Yang, F. (2020) Large-scale Identification of N-linked Intact Glycopeptides in Human Serum using HILIC Enrichment and Spectral Library Search. Mol Cell Proteomics 19, 672–689

51. Zhu, J., Lin, Y. H., Dingess, K. A., Mank, M., Stahl, B., and Heck, A. J. R. (2020) Quantitative Longitudinal Inventory of the N-Glycoproteome of Human Milk from a Single Donor Reveals the Highly Variable Repertoire and Dynamic Site-Specific Changes. J Proteome Res 19, 1941–1952

52. Leung, K. K., Wilson, G. M., Kirkemo, L. L., Riley, N. M., Coon, J. J., and Wells, J. A. (2020) Broad and thematic remodeling of the surfaceome and glycoproteome on isogenic cells transformed with driving proliferative oncogenes. Proc Natl Acad Sci U S A 117, 7764–7775

53. Li, J., Jia, L., Hao, Z., Xu, Y., Shen, J., Ma, C., Wu, J., Zhao, T., Zhi, Y., Li, P., Li, J., Zhu, B., and Sun, S. (2020) Site-specific N-glycoproteme Analysis Reveals Up-regulated Sialylation and Core Fucosylation during Transient Regeneration Loss in Neonatal Mouse Hearts. J Proteome Res

54. Medzihradszky, K. F., Kaasik, K., and Chalkley, R. J. (2015) Tissue-Specific Glycosylation at the Glycopeptide Level. Mol Cell Proteomics 14, 2103–2110

55. Liu, M. Q., Zeng, W. F., Fang, P., Cao, W. Q., Liu, C., Yan, G. Q., Zhang, Y., Peng, C., Wu, J. Q., Zhang, X. J., Tu, H. J., Chi, H., Sun, R. X., Cao, Y., Dong, M. Q., Jiang, B. Y., Huang, J. M., Shen, H. L., Wong, C. C. L., He, S. M., and Yang, P. Y. (2017) pGlyco 2.0 enables precision N-glycoproteomics with comprehensive quality control and one-step mass spectrometry for intact glycopeptide identification. Nat Commun 8, 438

56. Chalkley, R. J., Medzihradszky, K. F., Darula, Z., Pap, A., and Baker, P. R. (2020) The effectiveness of filtering glycopeptide peak list files for Y ions. Mol Omics 16, 147–155

57. Klein, J., and Zaia, J. (2020) Relative Retention Time Estimation Improves N-Glycopeptide Identifications by LC-MS/MS. J Proteome Res 19, 2113–2121

58. Yang, Y., Franc, V., and Heck, A. J. R. (2017) Glycoproteomics: A Balance between High-Throughput and In-Depth Analysis. Trends Biotechnol 35, 598–609

59. Wandall, H. H., Pizette, S., Pedersen, J. W., Eichert, H., Levery, S. B., Mandel, U., Cohen, S. M., and Clausen, H. (2005) Egghead and brainiac are essential for glycosphingolipid biosynthesis in vivo. J Biol Chem 280, 4858–4863

60. Langeveld, M., and Aerts, J. M. (2009) Glycosphingolipids and insulin resistance. Prog Lipid Res 48, 196–205

61. Ruvolo, P. P. (2019) Galectins as regulators of cell survival in the leukemia niche. Adv Biol Regul 71, 41–54

62. Mendez-Huergo, S. P., Hockl, P. F., Stupirski, J. C., Maller, S. M., Morosi, L. G., Pinto, N. A., Beron, A. M., Musuruana, J. L., Nasswetter, G. G., Cavallasca, J. A., and Rabinovich, G. A. (2018) Clinical Relevance of Galectin-1 and Galectin-3 in Rheumatoid Arthritis Patients: Differential Regulation and Correlation With Disease Activity. Front Immunol 9, 3057

63. Boscher, C., Dennis, J. W., and Nabi, I. R. (2011) Glycosylation, galectins and cellular signaling. Curr Opin Cell Biol 23, 383–392

64. Ahmad, N., Gabius, H. J., Andre, S., Kaltner, H., Sabesan, S., Roy, R., Liu, B., Macaluso, F., and Brewer, C. F. (2004) Galectin-3 precipitates as a pentamer with synthetic multivalent carbohydrates and forms heterogeneous cross-linked complexes. J Biol Chem 279, 10841–10847

65. Rancourt, A., Dufresne, S. S., St-Pierre, G., Levesque, J. C., Nakamura, H., Kikuchi, Y., Satoh, M. S., Frenette, J., and Sato, S. (2018) Galectin-3 and N-acetylglucosamine promote myogenesis and improve skeletal muscle function in the mdx model of Duchenne muscular dystrophy. FASEB J, fj201701151RRR

66. Kamili, N. A., Arthur, C. M., Gerner-Smidt, C., Tafesse, E., Blenda, A., Dias-Baruffi, M., and Stowell, S. R. (2016) Key regulators of galectin-glycan interactions. Proteomics 16, 3111–3125

67. Hughes, R. C. (1999) Secretion of the galectin family of mammalian carbohydrate-binding proteins. Biochim Biophys Acta 1473, 172–185

68. Furtak, V., Hatcher, F., and Ochieng, J. (2001) Galectin-3 mediates the endocytosis of beta-1 integrins by breast carcinoma cells. Biochem Biophys Res Commun 289, 845–850

69. Chauhan, S., Kumar, S., Jain, A., Ponpuak, M., Mudd, M. H., Kimura, T., Choi, S. W., Peters, R., Mandell, M., Bruun, J. A., Johansen, T., and Deretic, V. (2016) TRIMs and Galectins Globally Cooperate and TRIM16 and Galectin-3 Co-direct Autophagy in Endomembrane Damage Homeostasis. Dev Cell 39, 13–27

70. Jia, J., Abudu, Y. P., Claude-Taupin, A., Gu, Y., Kumar, S., Choi, S. W., Peters, R., Mudd, M. H., Allers, L., Salemi, M., Phinney, B., Johansen, T., and Deretic, V. (2018) Galectins Control mTOR in Response to Endomembrane Damage. Mol Cell 70, 120–135 e128

71. Gordon, B. S., Kelleher, A. R., and Kimball, S. R. (2013) Regulation of muscle protein synthesis and the effects of catabolic states. Int J Biochem Cell Biol 45, 2147–2157

